# *Entamoeba histolytica* adaption to auranofin: a phenotypic and multi-omics characterization

**DOI:** 10.1101/2021.06.23.449586

**Authors:** Yana Shaulov, Lotem Sarid, Meirav Trebicz-Geffen, Serge Ankri

## Abstract

Auranofin (AF), an antirheumatic agent, targets mammalian thioredoxin reductase (TrxR), an important enzyme controlling redox homeostasisis, AF is also very effective against a diversity of pathogenic bacteria and protozoan parasites. Here, we report about the resistance of the parasite *Entamoeba histolytica* to 2 μM of AF that has been acquired by gradual exposure of the parasite to increasing amount of the drug. AF adapted *E.histolytica* trophozoites (AFAT) has an impaired growth, cytopathic activity and they are more sensitive to oxidative stress (OS), nitrosative stress (NS) and metronidazole (MTZ) than wild type (WT) trophozoites. Integrated transcriptomics and redoxomics analyses showed that many upregulated genes in AFAT, including genes encoding for dehydrogenase and cytoskeletal proteins, have their product oxidized in wild type trophozoites exposed to AF (acute AF trophozoites) but not in AFAT. We also showed that the level of reactive oxygen species (ROS) and oxidized proteins (OXs) in AFAT is lower than that of acute AF trophozoites. Overexpression of *E.histolytica* TrxR (EhTrxR) did not protect the parasite against AF which suggests that EhTrxR is not central is the mechanism of adaptation to AF.

## 1. Introduction

Amoebiasis is caused by the protozoan parasite *Entamoeba histolytica*. This disease is a significant hazard in underdeveloped countries with low socioeconomic status and poor sanitation. It is assessed that amoebiasis accounted for 55,500 deaths and 2.237 million disability-adjusted life years (the sum of years of life lost and years lived with disability) in 2010 [1]. Amoebiasis has also been diagnosed in tourists from developed countries who return from vacation in endemic regions. Inflammation of the large intestine and liver abscess represent the main clinical manifestations of amoebiasis. Amoebiasis is caused by the ingestion of food contaminated with cysts, the infective form of the parasite. Following excystation, the trophozoites migrate to the large intestine resulting in either asymptomatic colonization (90% of all infections) or causing bloody diarrhea. Metronidazole (MNZ) is the drug currently used for invasive amoebiasis [2]. Inside the parasite, MNZ is reduced through the action of thioredoxin reductase (TrxR) to a nitro radical anion or to a nitrosomidazole. This nitro group reduces O_2_ leading to the formation of cytotoxic reactive oxygen species (ROS) inside the parasite. The nitroso imidazole can also modify cysteine containing proteins such as thioredoxin (Trx) leading to their inactivation [3]. Common side effects related to MNZ are many and they include dizziness, heartburn, stomach cramps, trouble sleeping and weight loss [4–6]. Treatment with MNZ is usually very effective but resistance to this drug has been reported in various bacteria [7,8] and protozoan parasites [9–11]. To address these drawbacks, new alternatives to MNZ have been initiated and AF has emerged as one of the most potent anti-protozoan parasites drugs. Initially, AF is a gold containing compound that has been developed in the seventies for the treatment of rheumatoid arthritis [12]. Its mechanism of action as an antiarthritic gold drug remained controversial but it is assumed that it works by inhibiting the activity of TrxR, a crucial enzyme involved in the maintenance of the redox homeostasis in the cell [13]. AF is also a potent anticancer agent [14] and it has been found to be very efficient against a number of pathogens including *Mycobacterium abscessus* [15], *Clostridium difficile* [16,17], vancomycin-resistant enterococci [18,19], and some additional multidrug resistant bacteria [20]. Auranofin is also very efficient against parasites including the trematode *Schistosoma mansoni* [21,22] and protozoan parasites including *Trichomonas vaginalis* [23], *Giardia lamblia* [24] and *E.histolytica* [25]. The mode of action of AF in protozoan parasites is not completely understood although it is assumed that TrxR is the main target of AF in *E.histolytica* [25] [24]. In *G.lamblia*, this mechanism of action has been challenged by the significant TrxR activity present in trophozoites exposed to high concentrations of auranofin. Indeed, overexpression of TrxR in *G.lamblia* has no effect on the sensitivity of this parasite to AF [26]. AF can also target *E. histolytica* adenosine 5′-phosphosulfate kinase (EhAPSK), an essential enzyme in Entamoeba sulfolipid metabolism [27].We have recently shown that AF induced the formation of more than 500 oxidized proteins (OXs) in *E.histolytica* including some crucial enzymes for redox homeostasis and cytoskeletal proteins which are essential for *E.histolytica’s* cytoskeleton dependent virulence [28]. Knowledge about resistance to AF in bacteria and in protozoan is scanty. Recently, toxoplasma trophozoites resistant to AF (2 μM) have been successfully generated through chemical mutagenesis. The authors have identified point mutations in genes encoding redox-relevant proteins such as superoxide dismutase and ribonucleotide reductase. However, recapitulation of these mutations in the parasite did not confer resistance to AF suggesting that the mechanism of resistance is complex [29]. In this work, we are using a multi-omics approach to characterize an *E.histolytica* strain made resistant to AF (AFAT) by progressively adapting the parasite to 2 μM of AF. At this concentration, the drug is lethal to non-adapted parasites [25] [28].

## 2. Materials and Methods

### 2.1. E. histolytica culture

*E. histolytica* trophozoites, HM-1:IMSS strain (a kind gift of Prof. Samudrala Gourinath, Jawaharlal Nehru University, New Delhi, India) were grown under axenic condition at 37°C in TYI-S-33 medium prepared according to a previously reported protocol. [30]. The trophozoites were harvested during the logarithmic phase of growth by chilling the culture tubes at 4°C and pelleted by centrifugation at 500 × g for 5 min.

### 2.2. Adaptation of E.histolytica trophozoites to AF

The concentration of AF in *E.histolytica* trophozoites culture was progressively increased from 0 to 2 μM over a period of one month.

### 2.3. Growth Rate of WT trophozoites and AFAT

WT trophozoites or AFAT (5×10^4^/ml) were grown in a 14 ml tube in TYI-S-33 medium at 37 ◻ and the number of viable trophozoites were counted following 24, 48, and 72 h of culture by using the eosin dye exclusion method [31].

### 2.4. Viability of AFAT exposed to H_2_O_2_, paraquat, MTZ or GSNO

The viability of WT trophozoites and AFAT exposed to H_2_O_2_ (2.5 mM for 30 min), paraquat (2.5 mM for 24 hrs), MNZ (5 uM for 24 hrs) or GSNO (350 μM for 2 hrs) (Sigma-Aldrich, USA) was determined by the eosin dye exclusion method [31].

### 2.5. Measurement of cytopathic activity

Cytopathic activity was assayed against HeLa cells (ATCC) (using a previously described protocol [32].

### 2.6. RNA extraction

Total RNA was extracted from control trophozoites (WT) and AFAT using the TRI reagent kit, according to the manufacturer instructions (Sigma-Aldrich USA).

### 2.7. RNA sequencing (RNAseq): Library preparation and data generation

Six RNAseq libraries were produced according to manufacturer protocol (NEBNext UltraII Directional RNA Library Prep Kit, Illumina) using 800ng of total RNA. mRNAs pull-up was performed using Magnetic Isolation Module (NEB). All libraries were mixed into a single tube with equal molarity. The RNAseq data was generated on Illumina NextSeq500, 75 single-end read, high output mode (Illumina). Quality control was assessed using Fastqc (v0.11.5), reads were trimmed for adapters, low quality 3` and minimum length of 20 using CUTADAPT (v1.12). 83 bp single-end reads were aligned using STAR aligner (v2.6.0a), to a *Entamoeba histolytica* reference genome (Entamoeba_histolytica.JCVI-ESG2-1.0.dna.toplevel.fa) and annotation file (Entamoeba_histolytica.JCVI-ESG2-1.0.46.gff3) both downloaded from ENSEMBL (strain HM-1:IMSS, imported from the AmoebaDB (https://amoebadb.org/amoeba/app)). The number of reads per gene was counted using Htseq-count (v0.9.1) (parameters: −t CDS −i ID −m intersection-nonempty −s reverse).

### 2.8. Descriptive analysis

The statistical analysis was preformed using DESeq2 R package (version 1.20.0) [33].

### 2.9. Differential expression analysis

Results of the statistical analysis, i.e. list of the differentially expressed genes (DEGs) (p-value adjusted (padj) <0.01) are provided in the DESeq2_results_with_anno.xls file (Table S1). Genes with a fold change >1.5 were taken into account for further bioinformatics analysis. Gene symbol and gene name identification was performed using PANTHER Classification System software (http://www.pantherdb.org/) [34].

### 2.10. Availability of data

RNA-Seq data have been deposited at the Gene Expression Omnibus (http://www.ncbi.nlm.nih.gov/geo) under the accession number GSE178520.

### 2.11. Construction of HA-tagged EhTrxR trophozoites

For the construction of pJST4-EhTrxR expression vector that was used to express HA-tagged EhTrxR in the parasite, EhTrxR was amplified from *E.histolytica’s* genomic DNA using the primers 5’EhTrxR_ KpnI (ggtaccatgagtaatattcatgatg) and 3’EhTrxR_BamHI (ggatccatgagtttgaagcc). The resulting PCR product was cloned into the pGEM-T Easy vector system (Promega) and then digested with the restriction enzymes, KpnI and BamHI. The digested DNA insert was subcloned into the *E. histolytica* expression vector pJST4 which had been previously linearized with KpnI and BamHI. The pJST4 expression vector contains a tandem affinity purification tag for use in protein purification and identification [35]. This CHH-tag contains the calmodulin binding protein, hemagglutinin (HA), and histidine (His) residues and its expression is driven by an actin promoter.

### 2.12. Immunodetection of (HA)-tagged EhTrxR

*E. histolytica* control and HA-tagged EhTrxR trophozoites cytosolic proteins (40 μg) were prepared according to a published method [36] and resolved on a 10% SDS-PAGE in SDS-PAGE running buffer (25 mM Tris, 192 mM glycine, 0.1% SDS). The resultant protein bands were visualized after staining with Ponceau-S (Sigma-Aldrich, USA). Next, proteins were electrotransferred in protein transfer buffer (25mM Tris, 192mM glycine, 20% methanol, pH 8.3) to nitrocellulose membranes (Whatman, Protran BA83). The blots were first blocked using 3% skim milk and then probed with 1:500 mouse monoclonal HA antibody clone 12CA5 (a kind gift from Prof. Ami Aronheim) for 16 h at 4 °C. After incubation with the primary antibody, the blots were incubated with 1:5000 secondary antibody for one hour at RT (Jackson ImmunoResearch), and then developed using enhanced chemiluminescence (Bio RAD).

### 2.13. Viability assay

*E. histolytica* trophozoites control and EhTrxR overexpressing trophozoites (2.5×10^4^) were cultivated or not in presence of 2μM AF for 24 hours. The cells were harvested at 400×g for 5 min, stained with Propidium iodide (1 μg/mL) and analyzed by flow cytometry. Flow cytometry was performed using Cyan ADP (Dako®) and data from 10,000 cells were collected for each condition.

### 2.14. Detection of ROS

WT trophozoites, AFAT, and WT trophozoites that were cultivated with AF (2 μM) for 24 hours (WT+AF acute) were incubated with 0.4mM H2DCFDA for 15 min at 37°C. The trophozoites were washed twice with PBS, and the level of oxidation was analyzed by flow cytometry. Flow cytometry was performed using Cyan ADP (Dako®) and data from 10,000 cells were collected for each condition.

### 2.15. Detection of OX proteins by RAC (OX-RAC)

The detection of OXs by OX-RAC was performed using a previously described protocol [31]. Captured proteins were eluted with 30 μl elution buffer which contained 10 mM HEPES, 0.1 mM EDTA, 0.01 mM neocuproine, 0.1% sodium dodecyl sulfate (SDS) and 100 mM 2-mercaptoethanol for 20 min at room temperature. Proteins in a 10 μl aliquot of each eluent were resolved on a 12.5% SDS–polyacrylamide gel electrophoresis (PAGE) gel. Proteins in each gel were detected by silver staining (Pierce Silver Stain), and each gel slice was independently analyzed by MS. A protein was considered to be oxidized when its relative amount in the DTT-treated lysates was significantly less than that in the DTT-untreated lysates (*p* < 0.05 according to the results of an unpaired *t*-test).

### 2.16. In gel proteolysis and MS analysis

In gel proteolysis by trypsin and analysis by liquid chromatography– tandem mass spectrometry on the Q Exactive Plus (Thermo Fisher) and data analysis with MaxQuant 1.5.2.8 [37] and the UniProt database as the reference were done using a previously described protocol [31]. The data was quantified by LF analysis using the same software. The identifications are filtered for proteins identified with a false discovery rate of <0.01, and at least two identified peptides in the project. The intensities are presented as raw intensities without normalization and as label-free quantification with normalization, both presented as log2 intensities.

### 2.17. Classification of OXs according to their protein class

The OXs were classified according to their protein class using PANTHER Classification System software (http://www.pantherdb.org/) [34].

### 2.18. Immunofluorescence microscopy analysis

The formation of F-actin in WT trophozoites, acute AF trophozoites and AFAT was determined as described previously [28].

## 3. Results

### 3.1. Generation of AFAT and phenotypic characterization

Resistance to 2 μM of AF in the parasite *T.gondi* can be generated by exposure of the parasite to the mutagenic compound N-Ethyl-N-nitrosourea [29]. To our knowledge, the development by natural selection of parasites resistant to AF has never been attempted. In order to fill this knowledge gap, we adapted *E.histolytica* trophozoites to AF by progressively increasing the drug concentration over a period of one month to 2 μM. To check whether adaptation to AF has an effect on the growth of the parasite, we compared the doubling time of AF adapted trophozoites (AFAT) with that of wild type (WT) trophozoites. We found that the doubling time of AFAT (13 ±0.32 hours) was significantly higher than the doubling time of WT trophozoites (10.6 ± 0.24 hours). The response of AFAT to OS was tested by exposing them to H_2_O_2_ (2.5 mM for 30 min) or to paraquat (2.5 mM for 12 hours). We observed that AFAT are significantly more sensitive to H_2_O_2_ or to paraquat than WT trophozoites (Fig 1A). We also checked the resistance of AFAT to MNZ (5 μM for 24 hours) and we found that AFAT are significantly more sensitive to MNZ than WT trophozoites. (Fig. 1A). The sensitivity of AFAT to nitrosative stress (NS) has been tested by exposing them to the NO donor S-nitrosoglutathion (GSNO) (350 μM for 2 hours). We observed that AFAT are much more sensitive to NS than WT trophozoites (Fig 1A). The ability of AFAT to destroy a monolayer of mammalian cells (cytopathic activity) was also determined (Fig 1B). We observed that AFAT have their cytopathic activity impaired compared to that of WT trophozoites. Overall, these results indicate that for *E.histolytica* trophozoites, adaptation to AF comes at a cost to fitness.

**Figure 1.**
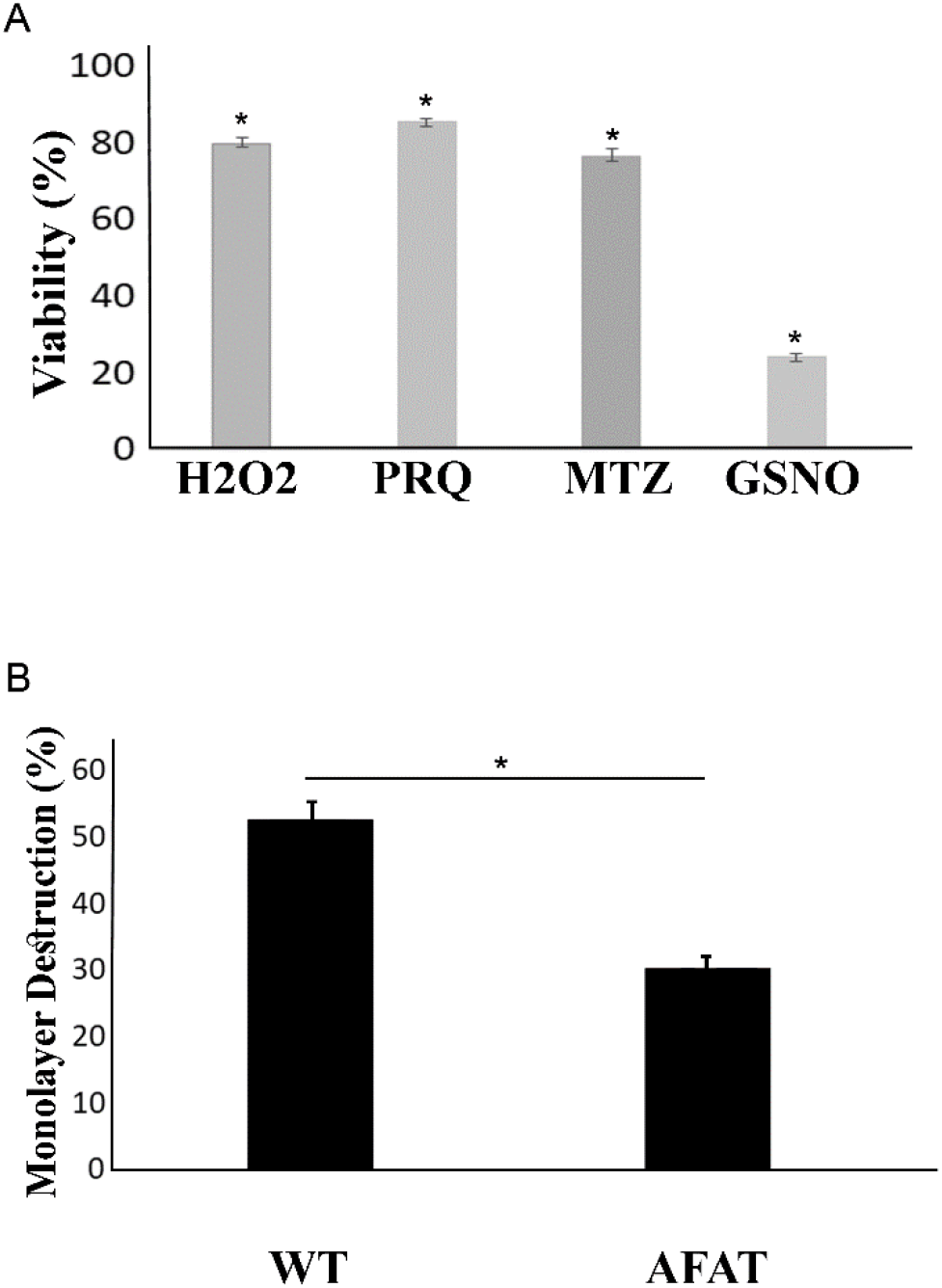
**A: Viability of AFAT exposed to H_2_O_2_, Paraquat, MTZ or to GSNO.** WT and AFAT were exposed to 2.5mM H_2_O_2_ for 30 minutes, 2.5mM paraquat (PRQ) and 5μM metronidazole (MTZ) for 24 hours or to 350μM GSNO for 2 hours. All experiments were done at 37 □. Data are expressed as the mean ± standard deviation of three independent experiments that were performed in triplicates. The graph represents the ratio percentage of viable amoebas compared to WT. The viability of AFAT exposed to H_2_O_2_, PRQ, MTZ or GSNO was significantly different (p < 0.05) from that of the WT according to the results of an unpaired Student’s t test. **B: Cytopathic activity of AFAT.** Data are displayed as the mean ± standard deviation of four independent experiments that were performed in triplicates. The cytopathic activity of AFAT was significantly different (p <0.05) from that of the WT according to the results of an unpaired Student’s t test.

### 3.2. Transcriptomics of AFAT

We used RNA sequencing (RNA-seq) to investigate the molecular basis of the adaptation of *E.histolytica* to AF. Transcriptomics of WT trophozoites vs AFAT was compared. Our comparisons revealed that adaptation to AF has a profound impact on the *E. histolytica* transcriptome with more than 500 upregulated and downregulated genes (Table S1).

### 3.3. Gene categories modulated in AFAT

The differentially regulated genes in AFAT vs WT trophozoites were classified according to the protein class which they encode using the Protein ANalysis THrough Evolutionary Relationship (PANTHER) sequence classification tool (version 14.1)[38]. The categories for functional classification of genes upregulated in AFAT (using the subset of protein class) is shown in Fig 2A. The most abundant class of gene encoded protein-binding activity modulator (PC00095) such as AIG1-type G domain-containing protein (EHI_176590), metabolite interconversion enzyme (PC00262) such as Lecithin:cholesterol acyltransferase (EHI_065250), protein modifying enzyme (PC00260) such as Leucine rich repeat / protein phosphatase 2C domain containing protein (EHI_178020) and cytoskeletal protein (PC00085) such as F-actin-capping protein subunit beta (EHI_134490). Of the upregulated genes in AFAT, genes that are encoding for actin or for actin-binding cytoskeletal proteins such as actin (EHI_107290) or EHI_172960 (Actin-related protein 2/3 complex subunit 3), dehydrogenase (PC00092) such as NAD (FAD)-dependent dehydrogenase (EHI_099700) or Aldehyde-alcohol dehydrogenase 2 (EHI_024240), guanyl-nucleotide exchange factor such as Ras guanine nucleotide exchange factor (EHI_023270) or Rho guanine nucleotide exchange factor (EHI_005910) are significantly enriched according to the PANTHER statistical overrepresentation test (Fig 2B).

**Figure 2:**
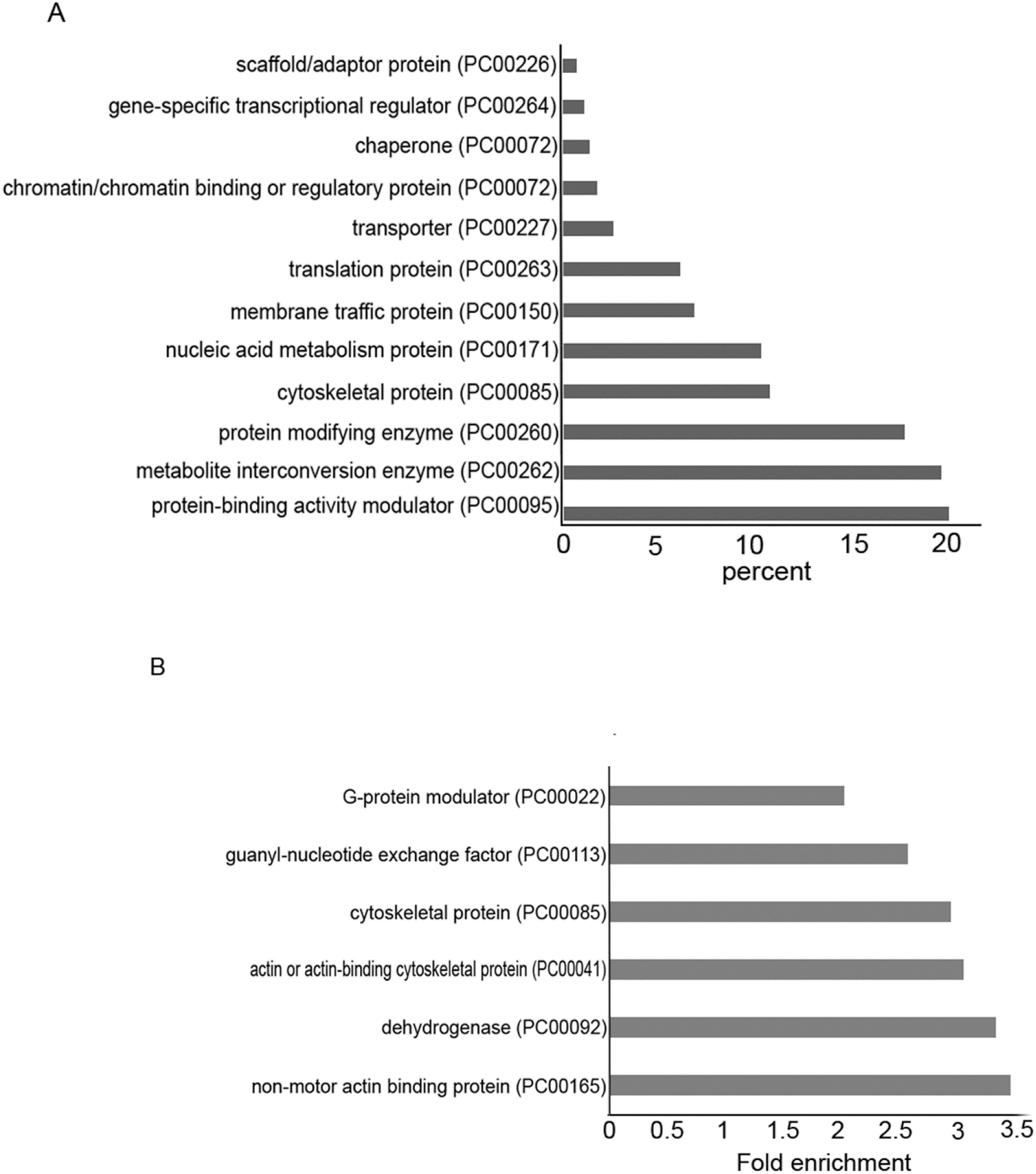
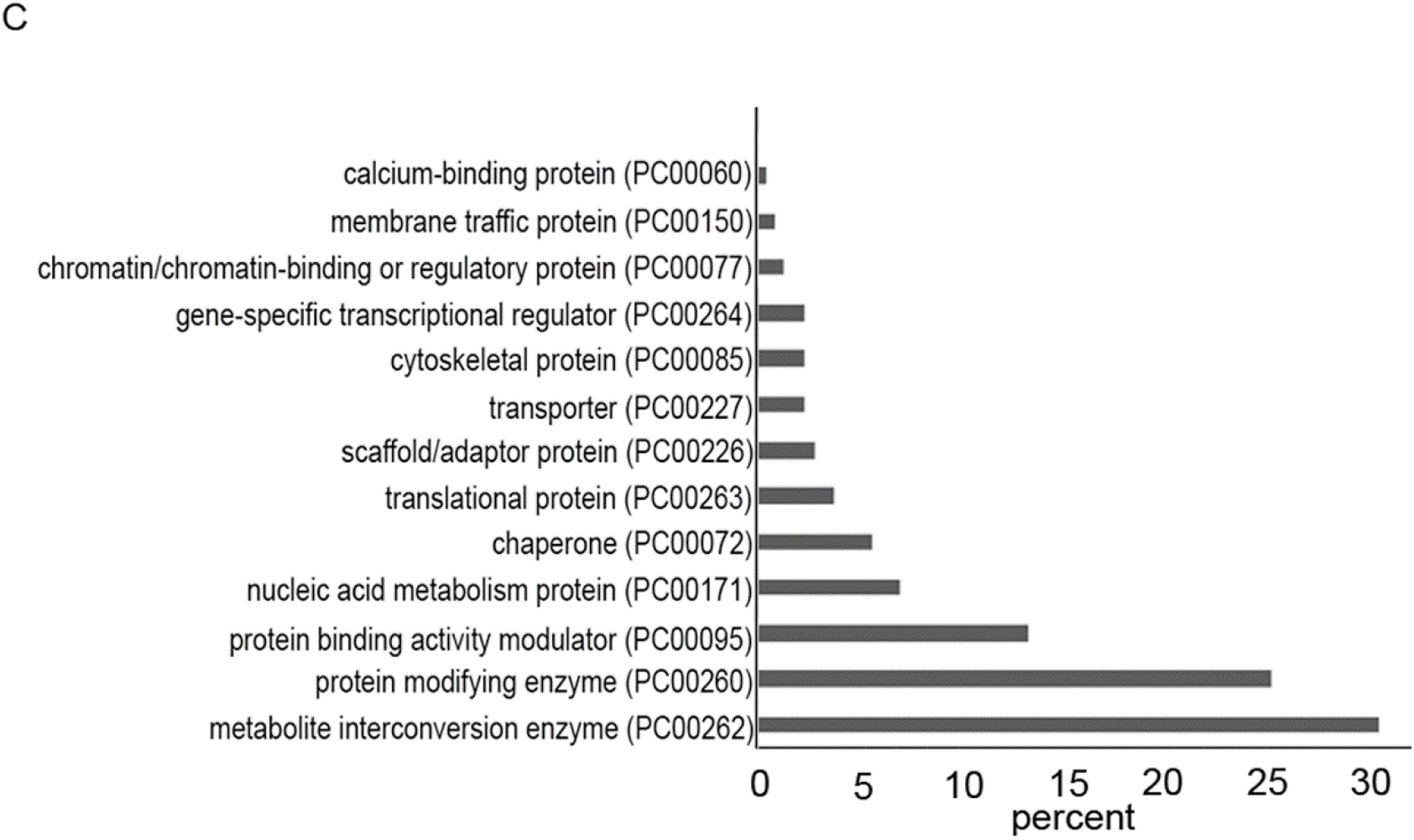
A. PANTHER sequence classification of genes upregulated in AFAT. B. PANTHER statistical overrepresentation test of upregulated genes in AFAT C. PANTHER sequence classification of genes downregulated in AFAT.

The categories for functional classification of genes downregulated in AFAT (using the subset of protein class) is shown in Fig 2C. The most abundant class of gene encoded proteins are metabolite interconversion enzyme (PC00262) such as alpha-amylase (EHI_152880), protein modifying enzyme (PC00260) such as Gal/GalNAc lectin Igl2 (EHI_183000) and protein-binding activity modulator (PC00095) such as guanylate binding protein (EHI_175080). Of the downregulated genes in AFAT, no enrichment of a specific biological process was detected according to the PANTHER statistical overrepresentation test.

### 3.4. Redoxomics of AFAT

We have previously detected by OX-RAC 583 OXs in acute AF trophozoites [28]. Here, we also used OX-RAC to detect OXs in the lysate of AFAT (Fig 3A). The purification of OXs by OX-RAC has been previously described in detail [31]. We identified 96 OXs in AFAT (Table S2) that were classified using PANTHER sequence classification tool. The most abundant OXs families belong to metabolite interconversion enzyme (PC00262) such as Purine nucleoside phosphorylase (EHI_200080), protein modifying enzyme (PC00260) such as NEDD8-activating enzyme E1 (EHI_098550), chaperone (PC00072) such as Peptidylprolyl isomerase (EHI_044850) and Protein-binding activity modulator (PC00095) such as glucosidase 2 subunit beta (EHI_135420) (Fig 3B).

**Figure 3:**
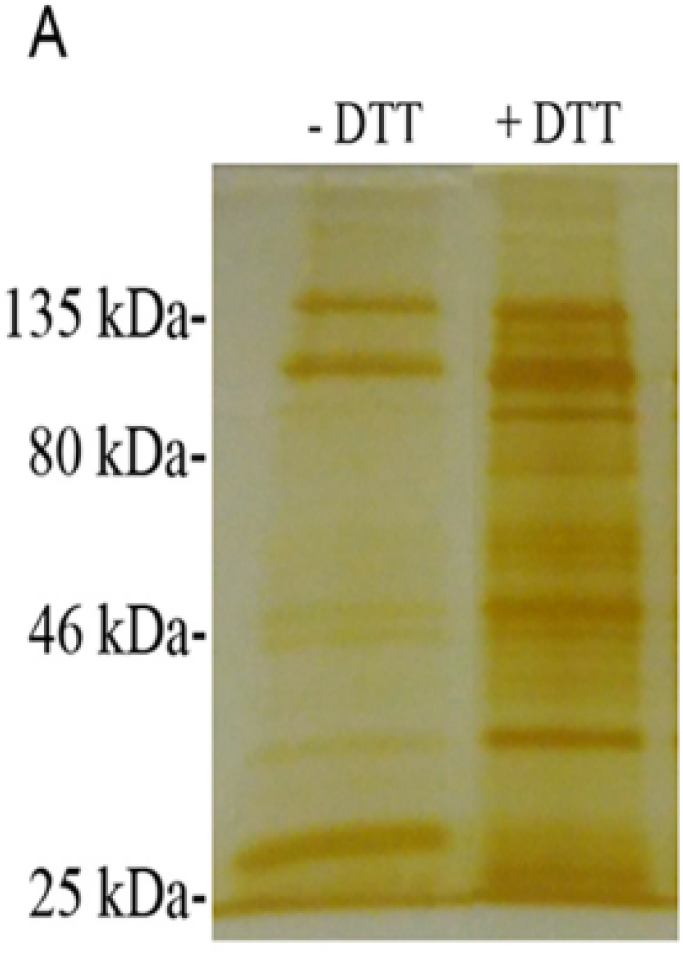
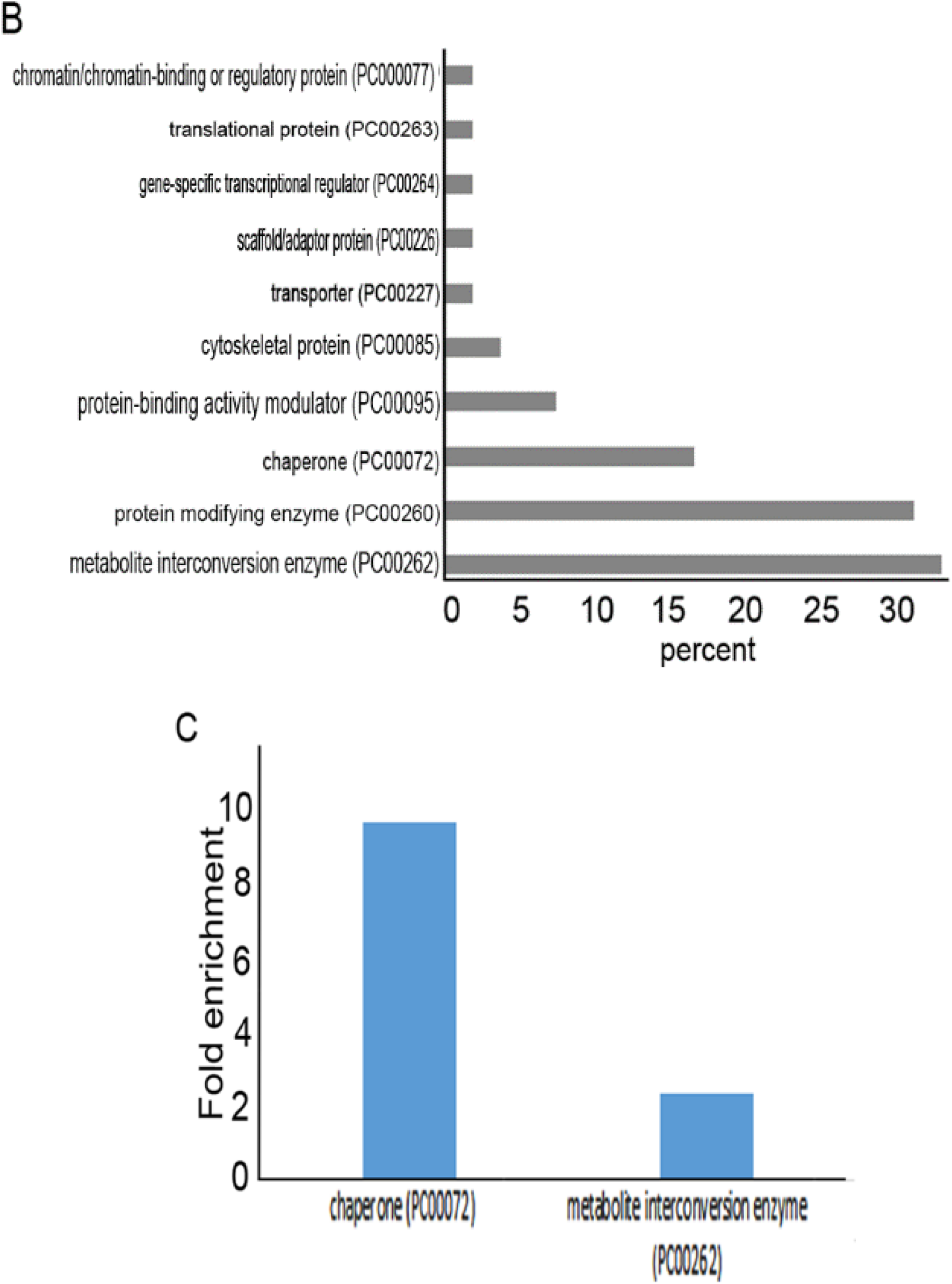
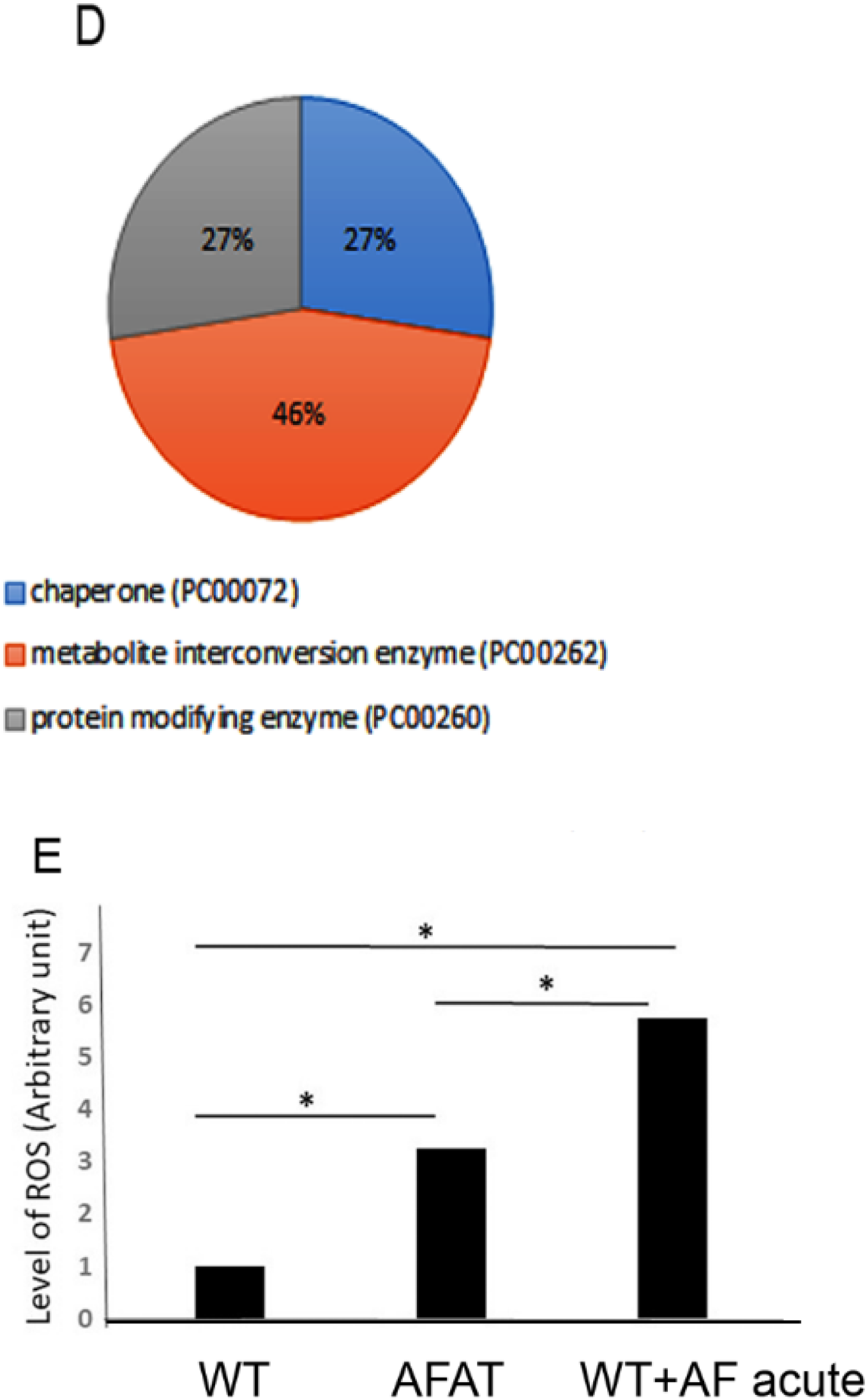
Detection of oxidized proteins by resin-assisted capture (OX-RAC) analysis of AFAT. A. Silver staining of OXs. OXs in the AFAT lysates were subjected to RAC in the presence of 10 mM DTT (+DTT) or the absence of DTT (−DTT). B. Protein ANalysis THrough Evolutionary Relationships (PANTHER) sequence classification of the OXs identified in AFAT. C. PANTHER statistical overrepresentation test of the OXs identified in AFAT D. PANTHER sequence classification of the 17 OXs common between trophozoites exposed to an acute AF treatment [28] and AFAT. E. Level of ROS in AFAT and acute AF trophozoites. WT trophozoites, AFAT, and WT trophozoites that were cultivated with AF (2 μM) for 24 hours (WT+AF acute) were incubated with 0.4mM H2DCFDA for 15 minutes at 37°C. The trophozoites were washed twice with PBS, and the level of oxidation was analyzed by flow cytometry. Flow cytometry was performed using Cyan ADP (Dako®) and data from 10,000 cells were collected for each condition. Data are expressed as the mean ± standard deviation of three independent experiments. The level of ROS in AFAT was significantly different from that of the WT+AF acute according to the results of an unpaired Student’s t test (p value < 0.05).

Of the OXs in AFAT (table S2), chaperone (PC00072) such as HSP16 (EHI_125830) or Trx, (EHI_110350) and metabolite interconversion enzyme (PC00262) such as are Aminotran_5 domain-containing protein EhnifS (EHI_136380) or alpha-amylase EHI_152880 are significantly enriched according to the PANTHER statistical overrepresentation test (Fig 3C).

Seventeen OXs are shared between acute AF trophozoites [28] and AFAT (Table S3). These common OXs belong to chaperone (PC00072), metabolite interconversion enzyme (PC00262) and protein modifying enzyme (PC00260) (Fig 3D).

The lower amount of OXs in AFAT compared to the amount of OXs in acute AF trophozoites [28] suggests that AFAT are less exposed to ROS. To test this hypothesis, we measured the level of ROS with dichloro-fluorescein (H2DCDFC) in acute AF trophozoites and in AFAT. We observed that the level of ROS in AFAT is significantly lower than in acute AF trophozoites (Fig 3E).

### 3.5. Comparison between transcriptomics and redoxomics of AFAT

We found that only two genes upregulated in AFAT (Gal/GalNAc lectin Igl1 EHI_006980, and SNF7 family protein EHI_077530) have their product oxidized (Table S3). None of the genes downregulated in AFAT have their product oxidized (Table S3).

### 3.6. Comparison between transcriptomics and redoxomics of acute AF trophozoites

We found that 77 genes up regulated in AFAT have their product oxidized in acute AF trophozoites [28] (Table S3). The most abundant OXs belong to metabolite interconversion enzyme (PC00262), protein-binding activity modulator (PC00095), protein modifying enzyme (PC00260) and cytoskeletal protein (PC00085) (Fig 4A). Of the upregulated genes in AFAT that have their product oxidized in acute AF trophozoites, genes that are encoding for dehydrogenase (PC00092) such as NAD(FAD)-dependent dehydrogenase (EHI_099700), oxydoreductase (PC00176) such as Pyruvate:ferredoxin oxidoreductase (EHI_051060) and metabolite interconversion enzymes (PC00262) such as isopentenyl phosphate kinase (EHI_178490) are significantly enriched according to the PANTHER statistical overrepresentation test (Fig 4B).

**Figure 4.**
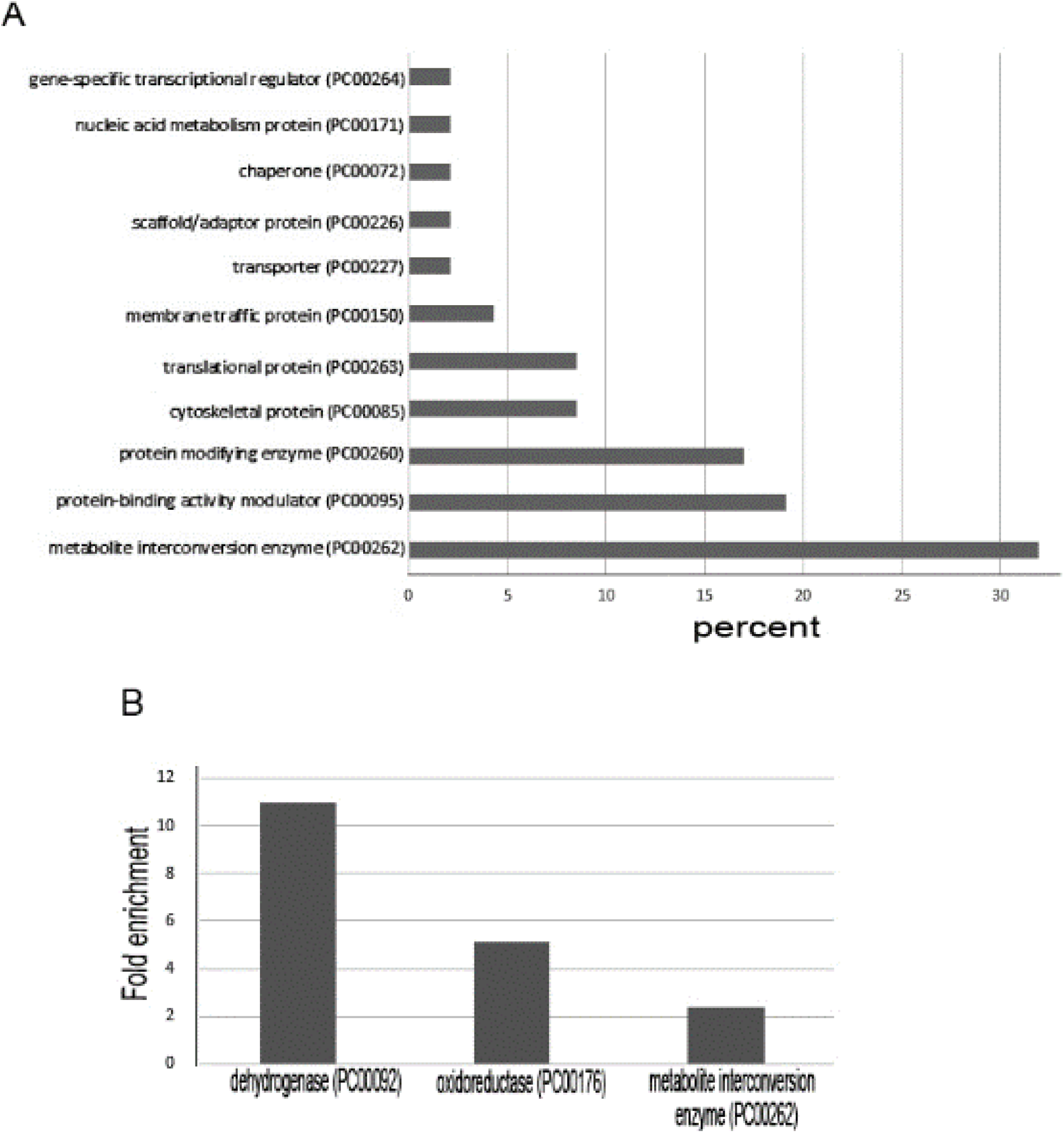
Comparison between transcriptomics and redoxomics of acute AF trophozoites. **A.** PANTHER sequence classification of genes upregulated in AFAT that have their product oxidized in acute AF trophozoites. **B.** PANTHER statistical overrepresentation test of upregulated genes in AFAT that have their product oxidized in acute AF trophozoites.

Eight genes that are downregulated in AFAT have their product oxidized in acute AF trophozoites [28] (Table S3). These OXs are the uncharacterized proteins (EHI_008120, EHI_065710 and EHI_110780), Asparagine--tRNA ligase (EHI_126920), Cytosolic Fe-S cluster assembly factor NUBP1 (EHI_047750), ribonuclease (EHI_156310) and Flavodoxin-like domain-containing protein (EHI_096710).

### 3.7. Formation of F-actin in AFAT

We have previously shown that AF leads to the oxidation of cytoskeletal proteins and inhibits the formation of actin filaments (F-actin) [28]. In contrast, cytoskeletal proteins in AFAT are not significantly enriched among OXs according to the PANTHER statistical overrepresentation test (table S2). In order to confirm this observation, we looked at the level of F-actin in WT trophozoites, acute AF trophozoites and in AFAT by immunofluorescence microscopy with phalloidin, a molecule that binds selectively to F-actin [39]. As described previously [28], the intensity of the F-actin signal in acute AF trophozoites was significantly less than in WT trophozoites. In contrast, the F-actin signal was identical in WT trophozoites and AFAT (Fig 5A&B). These results confirm that the formation of F-actin is impaired in acute AF trophozoites [28] but is not impaired in AFAT.

**Figure 5.**
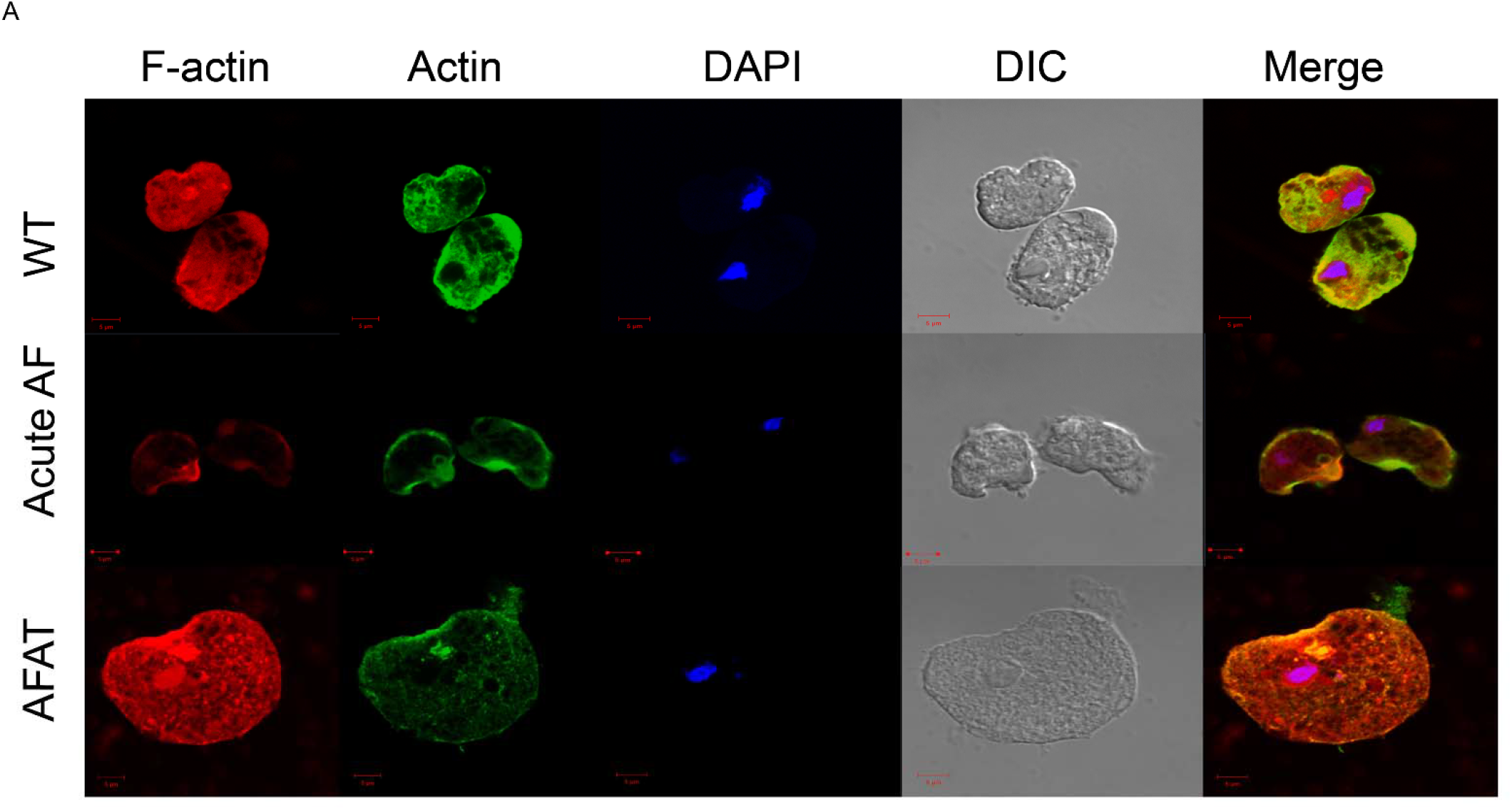
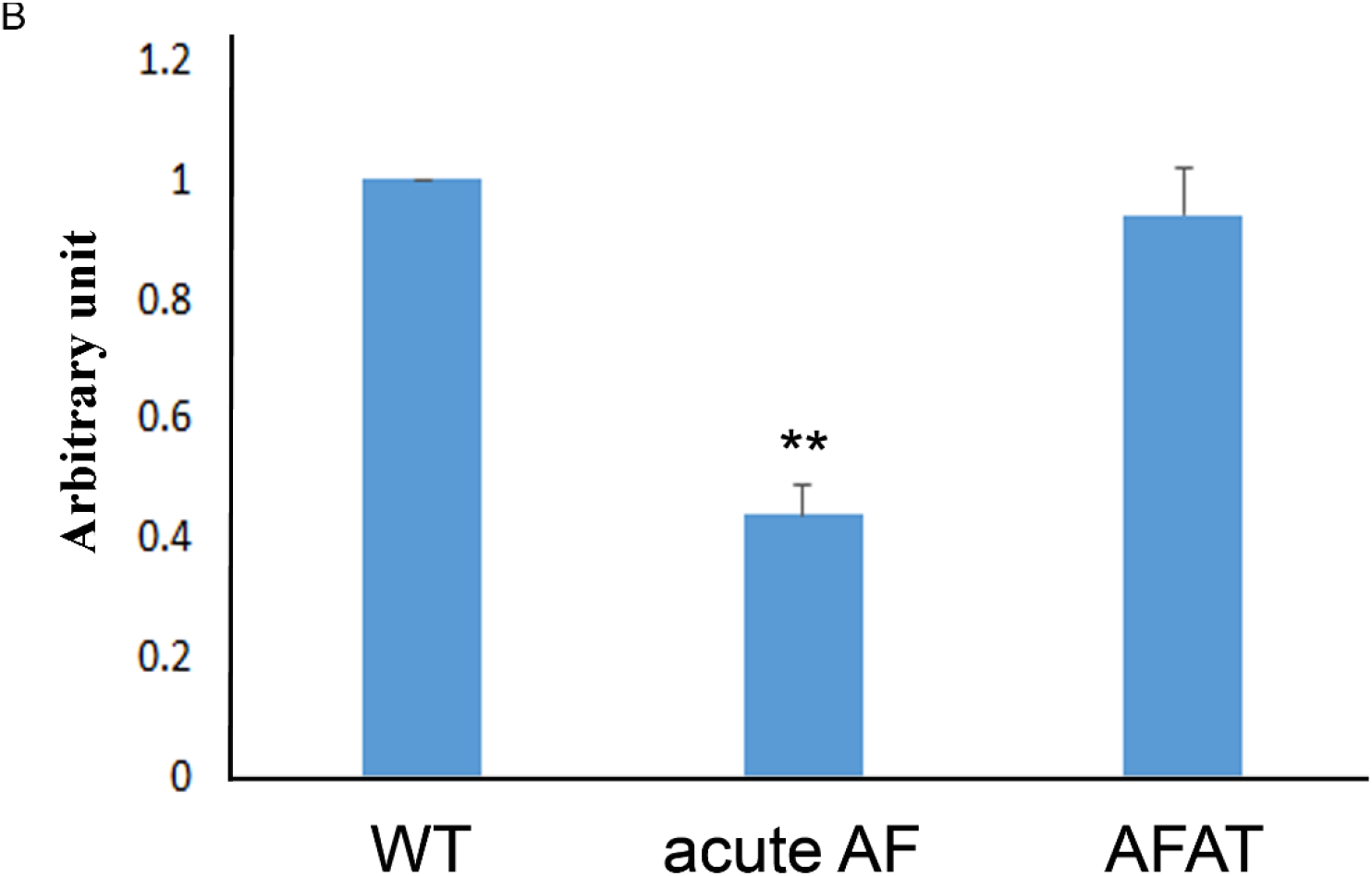
Formation of F-actin in WT trophozoites, acute AF trophozoites and AFAT. Confocal laser scanning microscopy of F-actin and total actin in WT trophozoites, acute AF trophozoites and AFAT. F-actin was detected using rhodamine-conjugated phalloidin. Total actin was detected using a primary actin antibody and a secondary Cy2-conjugated immunoglobulin G (IgG) antibody. The nuclei (blue) were stained by 4',6-diamidino-2-phenylindole (DAPI). Computer-assisted image overlay of the signal emitted by the actin antibody, phalloidin, and DAPI. Fluorescence quantification was performed using Fiji software [66] on 10 trophozoites and the F-actin signal was normalized to the total actin signal. The level of F-actin in WT was arbitrary defined as 1. Data are expressed as the mean ± standard deviation of two independent experiments. The level of F-actin in acute AF trophozoites was significantly different from that of WT and AFAT according to the results of an unpaired Student’s t test (p value < 0.01). No difference of F-actin level between WT and AFAT was observed according to the results on unpaired Student’s t test (p value > 0.05).

### 3.8. Overexpression of EhTrxR does not protect E.histolytica trophozoites against AF

Overexpression of TrxR in the parasite *Giardia lamblia*, has no effect on its resistance to AF [26]. In *E.histolytica*, Debnath *et al.* have found that AF inhibits the amebic TrxR, prevents its reduction and consequently enhances the sensitivity of trophozoites to reactive oxygen-mediated killing [40]. Our observations regarding the level of TrxR expression, which is the same in WT trophozoites and in AFAT (table S1) and the fact that Trxs are enriched OXs in AFAT, strongly suggest that *E.histolytica* TrxR is not central in the mechanism of adaptation of the parasite to AF. To test this hypothesis, we overexpressed EhTrxR in *E.histolytica* trophozoites. Overexpression of EhTrxR was confirmed by western blot and its level of expression in E.histolytica was proportional to the amount of G418 used for selection (Fig 6AB&C)[41]. Next, we determined the level of resistance to AF of HA-tagged EhTrxR trophozoites. We observed that the level of resistance to AF of HA-tagged EhTrxR trophozoites did not differ significantly from the level of resistance of control trophozoites (trophozoites transfected with pEhExGFP (a kind gift from Dr. Tomoyoshi Nozaki [42]) (Fig 6D). pEhExGFP allows the constitutive expression of the green fluorescent protein (GFP).

**Figure 6:**
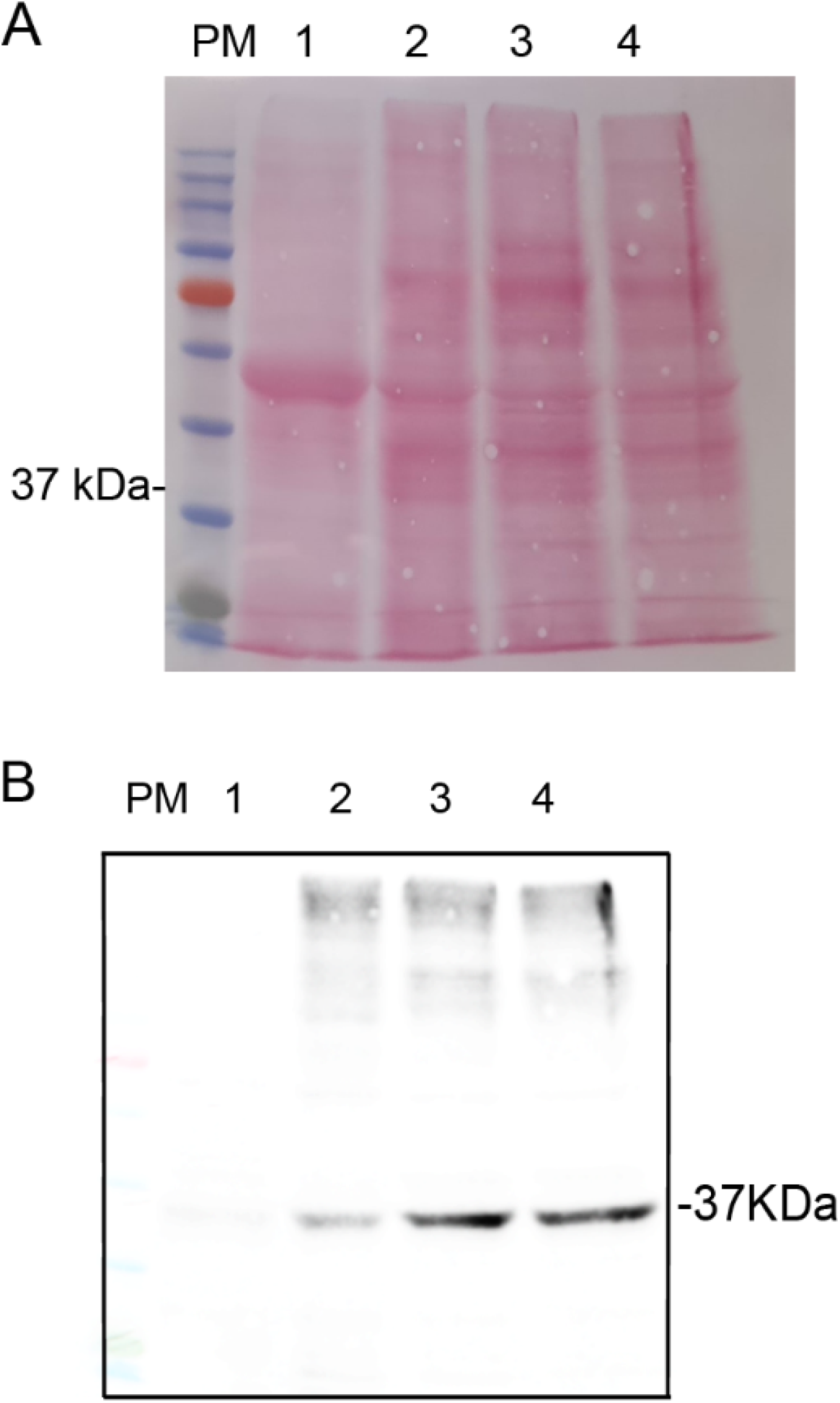
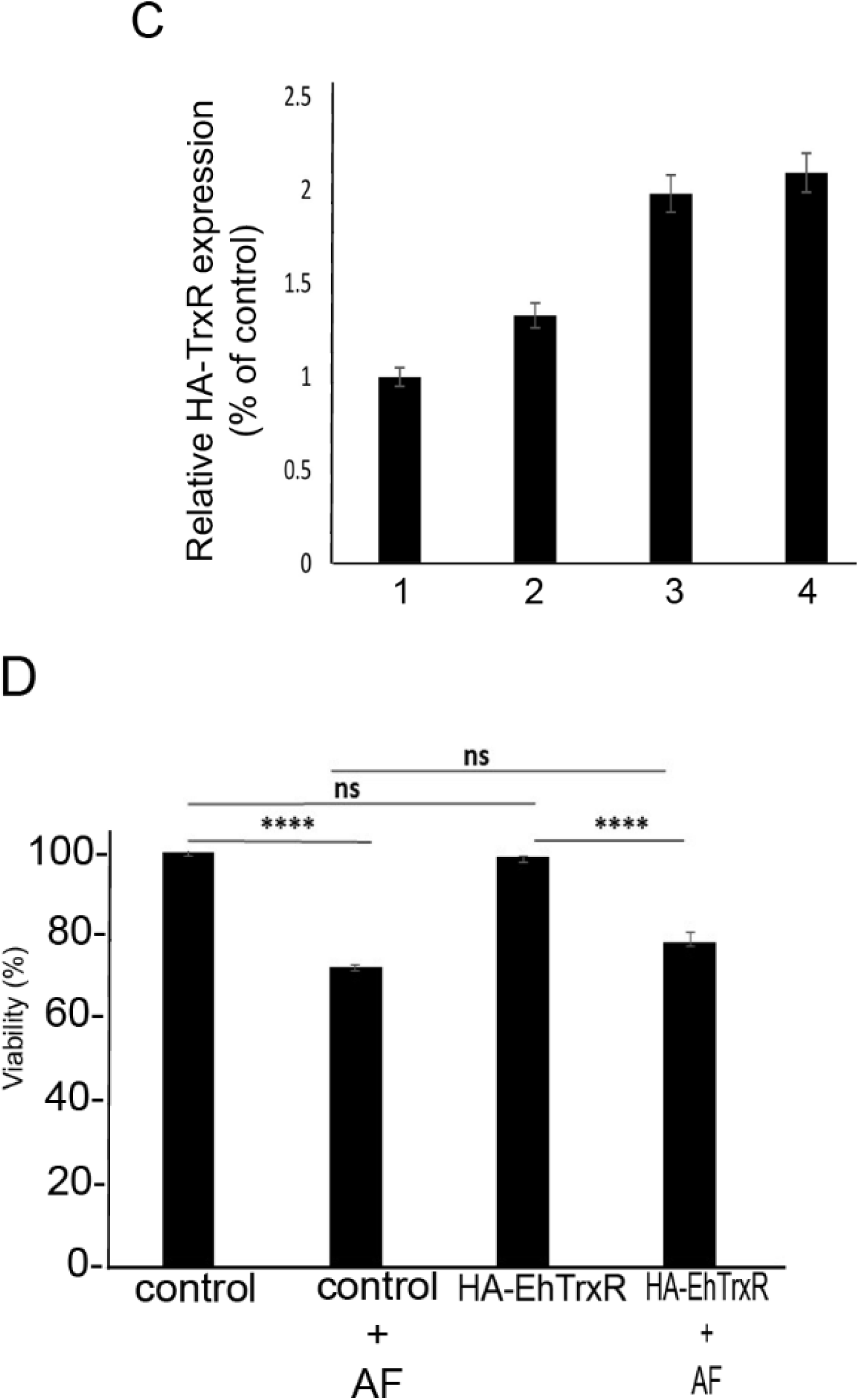
Western blot analysis of *E.histolytica* trophozoites that overexpress a hemagglutinin (HA)-tagged EhTrxR and viability assay. A. Ponceau staining of a nitrocellulose membrane containing cytosolic proteins (40 μg) separated by SDS PAGE of control trophozoites and of HA-tagged EhTrxR trophozoites cultivated in presence of increasing concentration of G418. B. Immunodetection of (HA)-tagged EHTrxR with an HA monoclonal antibody (1:500) in HA-tagged EhTrxR trophozoites cultivated in presence of increasing concentration of G418. C. Relative quantification of the HA EhTrxR signal following its normalization with the level of total protein in each well with the ImageJ software. Normalized values for control trophozoites were taken as 100%. These results are representative of two independent experiments. Legend: Protein molecular weight marker (PM). Control trophozoites (lane 1). HA-tagged EhTrxR trophozoites cultivated in presence of increasing concentration of G418 (lane 2: 6 μg/ml, lane 3: 30 μg/ml, lane 4:48 μg/ml). D. Viability assay. *E. histolytica* trophozoites control and HA-tagged EhTrxR trophozoites were cultivated in presence of 2μM AF for 24 hours. The cells were harvested at 400×g for 5 minutes, stained with Propidium iodide and analyzed by flow cytometry. Flow cytometry was performed using Cyan ADP (Dako®) and data from 10,000 cells were collected for each condition. Data are expressed as the mean ± standard deviation of three independent experiments that were done in triplicates. The viability of control trophozoites was defined as 100%. The viability of control trophozoites was not significantly different from that of the HA-tagged EhTrxR trophozoites according to the results of an unpaired Student’s t test (p value < 0.05).

## 4. Discussion

In our previous work, we have demonstrated that AF triggers OS inside *E.histolytica* trophozoites resulting in the oxidation of more than 500 proteins including many redox enzymes which are essential for maintaining intracellular levels of reactive oxygen species in the parasite [28,43,44]. Here, we have characterized *E.histolytica* trophozoites that have been adapted to 2 μM AF. Adaptation of *E.histolytica* to AF leads to the upregulation and downregulation of hundreds of genes, which suggests that the mechanism of adaptation is complex. Overexpression of a drug’s molecular target often increases drug resistance [45,46]. Consequently, we expected that *E.histolytica* TrxR (EhTrxR), the assumed main target of AF [25], would be one of the upregulated genes in AFAT. However, transcriptomics of AFAT indicates that this was not the case. Indeed, the overexpression of EhTrxR did not confer to *E.histolytica* resistance to AF. This information raises a question about why EhTrxR expression is not upregulated as a simple mechanism to resist AF. One possible answer is that, as for *Giardia lamblia*, TrxR is not the primary target of AF in *E.histolytica* [26]. This is supported by the absence of detection of EhTrxR among OXs in AFAT (this work) and acute AF trophozoites [28].

It is also possible that the fitness cost for *E.histolytica* to overexpress TrxR during adaptation to AF resistance is too high. EhTrxR has NAD(P)H dependent oxidase activity, which generates hydrogen peroxide (H_2_O_2_) from molecular oxygen leading to the formation of reactive species [47]. Therefore, it is possible that the production of H_2_O_2_ resulting from EhTrxR overexpression combined to OS triggered by AF [28] during the adaptation process cannot be tolerated by the parasite.

In this work, we found that only two genes upregulated in AFAT have their products oxidized in AFAT. In contrast, 77 genes upregulated in AFAT have their product oxidized in acute AF trophozoites [28]. The upregulation of these 77 genes in AFAT may be essential for the adaption of the parasite to AF by replacing their oxidized-inactivated-products by reduced-active proteins. The relevance of this mechanism for some of these 77 genes is discussed here:

Pyruvate:ferredoxin oxidoreductase (EHI_051060), NADP-dependent alcohol dehydrogenase (EHI_107210) and Fe-ADH domain-containing protein (EHI_198760) which encode for proteins involved in redox regulation. These redox enzymes depend on cysteine residues for their activity [48–50]. The oxidation of these cysteine residues impairs their activity [48,51].

Genes that encode protein-binding activity modulator like Ras guanine nucleotide exchange factor (EHI_035800), Rho guanine nucleotide exchange factor (EHI_005910) or Ras GTPase-activating protein (EHI_105250). These proteins have their product oxidized in acute AF trophozoites [28]. G proteins have been involved in key pathogenic processes of *E. histolytica* including vesicular trafficking and cytoskeleton regulation [52]. Redox regulation of G-proteins have been well documented [53] and their oxidation impairs *E.histolytica’s* motility [28].

Genes that encode protein-modifying enzymes like protein kinase domain-containing proteins (EHI_186820) (EHI_101280) and Protein kinase (EHI_188110) which are also oxidized in acute AF trophozoites [28]. Protein kinases have been associated with virulence and phagocytic activity of *E.histolytica* [54]. The redox regulation of protein kinases is well established [55] and it has been demonstrated that AF can directly inhibit protein kinase C by interacting with thiol groups present in the catalytic site [56].

Genes that encode actin or for actin-binding cytoskeletal proteins are upregulated in AFAT and oxidized in acute AF trophozoites [28]. In our previous work, we have shown that AF induces the oxidation of *E.histolytica* cytoskeletal proteins and consequently inhibits the formation of F-actin [28]. Consequently, it seems that the parasite upregulated the expression of actin-binding cytoskeletal proteins as a mechanism to adapt to AF by replacing oxidized cytoskeletal proteins that have been formed during the process of adaptation to AF. The low level of F-actin in acute AF trophozoites and the normal level of F-actin in AFAT (this work) support this hypothesis.

The fact that *E.histolytica* can adapt to AF illustrates the remarkable ability of *E.histolytica* to adapt to drugs [57,58] and environmental stresses [32,59]. The fitness cost paid by the parasite to adapt AF resembles collateral sensitivity (CS) that occurs when the acquisition of resistance to one antibiotic produces increased susceptibility to a second antibiotic [60]. AFAT are more sensitive to OS, paraquat, MNZ and GSNO than compared to WT trophozoites. Resistance to OS in *E.histolytica* involves the upregulation of 29-kDa peroxiredoxin [61] and iron-containing peroxide dismutase expression which is also involved in the resistance to MNZ [62] [63]. The level of expression of 29-kDa peroxiredoxin and iron-containing peroxide dismutase is globally the same in WT and in AFAT, which suggests that the sensitivity of AFAT to OS and MNZ is not caused by a reduced level of these redox enzymes expressions. We have discussed above that many oxidized proteins in AFAT have their level of expression upregulated. The fitness cost observed in AFAT may result from many reasons including the rerouting of protein synthesis toward oxidized proteins or substrate wasting that results from target overexpression [64]. In hydroxamic acid analog pan-histone deacetylase inhibitors resistant leukemia cells, overexpression of the target protein heat shock protein 90 (HSP90), revealed collateral sensitivity to the HSP90 inhibitor 17-N-allylamino-17-demethoxygeldanamycin [65].

## 5. Conclusions

We showed that *E.histolytica* trophozoites can be easily selected to resist toxic concentrations of AF in vitro. Adaptation to AF comes with a fitness cost for *E.histolytica* that includes a decreased growth rate and virulence and a sensitivity to OS, NS and MTZ. Overexpression of genes whose products are sensitive to AF-mediated oxidation may represent an important step in the adaptation process to AF and EhTrxR does not seem to be central for this process.

AF is FDA approved for the treatment of rheumatoid arthritis but it has not been yet used as an antimicrobial drug in the field. The ability of *E.histolytica* to adapt to amebicidal concentrations of AF raises an alarm for the future use of this drug as an antiamebic compound. Our omics data provides the basis for the development of strategies to limit the apparition of resistance against AF. One possible strategy suggested by our data is to promote dual antibiotic therapy (AF+MNZ) vs single AF therapy because adaptation to AF leads to more MNZ sensitivity in *E.histolytica*.

## Supplementary Materials

The following are available online at www.mdpi.com/xxx/s1, Figure S1: title, Table S1: title, Video S1: title.

## Author Contributions

Conceptualization: Y.S., S.A.; methodology, Y.S., S.A.; software, Y.S., S.A.; validation, Y.S, L.S., M.G, S.A.; formal analysis, Y.S, L.S., M.G, S.A.; investigation, Y.S, L.S., M.G, S.A.; resources, S.A.; data curation, Y.S., S.A.; writing—original draft preparation, Y.S., S.A‥; writing—review and editing, Y.S, L.S., M.G, S.A.; visualization, S.A‥; supervision, S.A.; project administration, S.A.; funding acquisition, S.A. All authors have read and agreed to the published version of the manuscript.

## Institutional Review Board Statement

In this section, please add the Institutional Review Board Statement and approval number for studies involving humans or animals. Please note that the Editorial Office might ask you for further information. Please add “The study was conducted according to the guidelines of the Declaration of Helsinki, and approved by the Institutional Review Board (or Ethics Committee) of NAME OF INSTITUTE (protocol code XXX and date of approval).” OR “Ethical review and approval were waived for this study, due to REASON (please provide a detailed justification).” OR “Not applicable.” for studies not involving humans or animals. You might also choose to exclude this statement if the study did not involve humans or animals.

## Informed Consent Statement

“Not applicable.”

## Acknowledgments

We thank the staff of the Microscopy Imaging facility, the genomics Core Facility Laboratory and the Smoler Proteomics Center at the Technion for their technical help.

