## Supplemental Table 2 legend for "*Entamoeba histolytica* adaption to auranofin: a phenotypic and multi-omics characterization"

| Proteins columns | |
|---|---|
| Parameter | Description |
| # AAs | Displays the length of the protein sequence. |
| # Decoy Proteins | Displays the number of the higher-ranked decoy or reverse proteins.<br>This column appears when the workflow includes the Protein FDR Validator node. |
| # Peptides | Displays the total number of distinct peptide sequences identified from all included searches. |
| # Peptides (by Search Engine) | Displays the number of distinct peptide sequences in the protein.<br>This column appears when the consensus workflow includes the Protein Scorer node. |
| # Protein Groups | Displays the total number of protein groups. |
| # Protein Unique Peptides | Displays the total number of peptides that are unique to a particular protein. |
| # PSMs | Displays the number of identified peptide spectrum matches identified from all included searches, including those redundantly identified. |
| # PSMs (by Search Engine) | Displays the number of identified peptide spectrum matches identified from all included searches, including those redundantly identified.<br>This column appears when the consensus workflow includes the Protein Scorer node. |
| # Razor Peptides | Displays the number of razor peptides (that is, peptides shared among multiple protein groups or proteins) used to quantify the protein when you use razor peptides for quantification.<br>This column appears when you set the Peptides to Use parameter of the Precursor Ions Quantifier node or the Reporter Ions Quantifier node to All or Unique + Razor. |
| # Unique Peptides | Displays the total number of distinct peptide sequences unique to the protein group. |
| Abundance Ratio Adj. P-Values | Displays the p-values adjusted by using the Benjamini-Hochberg correction for the false discovery rate. |
| Abundance Ratio P-Value | Displays the P value of the sample group calculated by running the Tukey HSD test (post hoc) after an analysis of variance (ANOVA) test. The P value is a number between 0 and 1. If the null hypothesis states that there is no difference between the sample groups for the variance tests, one of the following pertains:<br>• A low P value means there is a low probability that the null hypothesis is false; that is, the sample groups show a significant difference.<br>• A high P value means there is a high probability that the null hypothesis is true; that is, the sample groups do not show a significant difference. |
| Abundance Ratios | Displays abundance ratios as normal space values. |

| | |
|---|---|
| | This column appears when there are sample ratios defined in the analysis setup. |
| Abundance Ratios (by Bio. Rep.) | Displays the abundance ratios of the biological replicates as normal space values. |
| Abundance Ratios (log2) | Displays the abundance ratios as log2 values. |
| Abundance Ratios (log2) (by Bio. Rep.) | Displays the abundance ratios of the biological replicates as log2 values. |
| Abundances | Displays the abundance values of the samples before scaling and normalization. |
| Abundances (by Bio. Rep.) | Displays the abundance values of the biological replicates. |
| Abundances (by Bio. Rep.) Counts | Displays the number of abundance values used to calculate the abundances of the biological replicates. |
| Abundances (Grouped) | Displays the abundance values of the sample groups.<br>A grouped abundance value is calculated as the arithmetic mean of all the replicate abundance values within a sample group. You can specify the sample grouping on the Grouping and Quantification page when you set up an analysis.<br>This column appears when you group samples in the analysis setup, and there is at least one sample group consisting of at least two samples. |
| Abundances (Grouped) Counts | Displays the number of samples with detected abundance values used to calculate the abundance of the whole sample group. |
| Abundances (Grouped) Standard Errors [%] | Displays the standard error of the abundance values of the samples in a sample group, normalized to the group's median abundance. |
| Abundances (Normalized) | Displays the normalized abundances values of the samples.<br>This column appears when you set the Normalization Mode parameter of the Precursor Ions Quantifier node or the Reporter Ions Quantifier node to Total Peptide Amount or Specific Protein Amount. |
| Abundances (Scaled) | Displays the normalized and scaled abundance values of the samples.<br>This column appears when you set the Scaling Mode parameter of the Precursor Ions Quantifier node or the Reporter Ions Quantifier node to On All Average or On Controls Avg. |
| Abundances Counts | Displays the number of abundance values used to calculate the sample abundance. |
| Accession | Displays by default the unique identifier assigned to the protein by the FASTA database used to generate the report. |

| | |
|---|---|
| Biological Process | Displays the GO Slim categories of the protein’s biological processes.<br>This column appears when the consensus workflow includes the Protein Annotation node. |
| calc. pI | Displays the theoretically calculated isoelectric point for the protein, which is the pH at which a particular molecule carries no net electrical charge.<br>The amino acids that make up proteins can be positive, negative, neutral, or polar in nature, and together they give a protein its overall charge. At a pH below their isoelectric point, proteins carry a net positive charge; at a pH above their isoelectric point, they carry a net negative charge. Gel electrophoresis can then separate proteins according to their isoelectric point (overall charge) with a polyacrylamide gel, using a technique called isoelectric focusing. This technique uses a pH gradient to separate proteins and is the first step in two-dimensional gel polyacrylamide gel electrophoresis.<br>When you have searched the fractions resulting from isoelectric focusing, you can use the calc. pI value to estimate whether you might expect to find a particular protein in the given fraction. |
| Protein FDR Confidence | Displays the level of confidence for the identified protein as determined by the Protein FDR Validator node.<br>This column appears when the consensus workflow includes the Protein FDR Validator node in the consensus workflow. |
| Protein FDR Group Confidence | Displays the level of confidence for the identified protein group as determined by the Protein FDR Validator node.<br>This column appears when the consensus workflow includes the Protein FDR Validator node. |
| Master | Indicates whether the protein is the master protein of a protein group. For some peptides, a list of proteins might contain this peptide sequence, but none of them is a master protein. This situation can occur if the peptide contains isoleucine at a position where the master protein has leucine or vice versa. |
| Unique Sequence ID | Displays a numeric identifier unique to each protein. When you export the protein data from multiple searches and combine them during data processing in Python, R, or a similar program, you can use the unique sequence IDs to identify the duplicate protein sequences from different runs by fast integer comparison instead of slow sequence comparison. |
| Protein Group IDs | Displays the identification numbers of the reference protein groups. |
| Biological Process | Displays the GO Slim categories of the protein’s biological processes.<br>This column appears when the consensus workflow includes the Protein Annotation node. |
| Cellular Component | Displays the GO Slim categories of the protein’s cellular components.<br>This column appears when the consensus workflow includes the Protein Annotation node. |

| | |
|---|---|
| Checked | Indicates whether the item is selected. |
| Chromosome | Displays chromosome information from the Ensembl genome database.<br>This column appears when the consensus workflow includes the Protein Annotation node. |
| Coverage [%] | Displays the percentage of the protein sequences covered by identified peptides. |
| Coverage [%] (by Search Engine) | Displays the percentage of the protein sequence covered by identified peptides. |
| Description | Provides the name of the protein exclusive of the identifier that appears in the Accession column. This description appears in the table by default. |
| Ensembl Gene ID | Displays annotations from the Ensembl genome database.<br>This column appears when the consensus workflow includes the Protein Annotation node. |
| Entrez Gene ID | Displays the Entrez Gene database identification of the gene that the protein is derived from. If the gene is not stored in the Entrez Gene database, the value displayed is 0.<br>This column appears when the consensus workflow includes the Protein Annotation node. |
| Exp. q-value | Displays the q-values derived from the validation. The values must be greater than the thresholds set by the Protein FDR Validator node.<br>This column appears when the consensus workflow includes the Protein FDR Validator node. |
| FASTA Title Lines | Displays the FASTA title of the protein. |
| Found in Files | Displays the best confidence of the PSMs of the protein that the application identified in the files. The colors of the boxes represent the following:<br>•Green: high confidence<br>•Yellow: medium confidence<br>•Red: low confidence<br>Blue: found but unidentified PSM. Only the results from precursor ion<br>•quantification searches contain blue boxes.<br>This column appears only when you include the Data Distributions node in the consensus workflow and set its Show Found in Files parameter to True. |
| Found in Fractions | Displays the best confidence of the PSMs of the protein that the application identified in the fractions. The colors of the boxes represent the following:<br>•Green: high confidence<br>•Yellow: medium confidence<br>•Red: low confidence<br>Blue: found but unidentified PSM. Only the results from precursor ion<br>•quantification searches contain blue boxes. |

| | |
|---|---|
| | This column appears only when you include the Data Distributions node in the consensus workflow and set its Show Found in Fractions parameter to True. |
| Found in Sample Groups | Displays the best confidence of the PSMs of the protein that the application identified in the sample groups. The colors of the boxes represent the following:<br>•Green: high confidence<br>•Yellow: medium confidence<br>•Red: low confidence<br>Blue: found but unidentified PSM. Only the results from precursor ion<br>•quantification searches contain blue boxes.<br>This column appears only when you include the Data Distributions node in the consensus workflow and set its Show Found in Sample Groups parameter to True. |
| Found in Samples | Displays the best confidence of the PSMs of the protein that the application identified in the samples. The colors of the boxes represent the following:<br>•Green: high confidence<br>•Yellow: medium confidence<br>•Red: low confidence<br>Blue: found but unidentified PSM. Only the results from precursor ion<br>•quantification searches contain blue boxes.<br>This column appears only when you include the Data Distributions node in the consensus workflow and set its Show Found in Samples parameter to True. |
| Gene Symbol | Displays the official gene name that is used in publications. This information is taken from the second line of the General page of the ProteinCard page.<br>This column appears when the consensus workflow includes the Protein Annotation node. |
| GO Accessions | Displays the GO terms contained in the graph of the annotated GO term of a protein. When you move the cursor over the GO term, the application displays the annotated GO term and all ancestor terms.<br>This column appears when the consensus workflow includes the Protein Annotation node. |
| KEGG Pathway Accessions | Displays the accessions from the KEGG PATHWAY database.<br>This column appears on the Proteins page when you include the Protein Annotation node in the consensus workflow. |
| KEGG Pathways | Displays the descriptions from the KEGG PATHWAY database.<br>This column appears on the Proteins page when you include the Protein Annotation node in the consensus workflow. |
| Master | Indicates whether a protein is a master protein in a protein group. |

| | |
|---|---|
| Modifications | Displays the modifications identified in the protein consolidated from all PSMs. The column shows confidence value if the IMP-ptmRS node was used in the processing workflow. |
| Molecular Function | Displays the GO Slim categories of the protein's molecular functions as colored boxes. This column appears when the consensus workflow includes the Protein Annotation node. |
| MW [kDa] | Displays the calculated molecular weight of the protein. The application calculates the molecular weight without considering PTMs.<br>Separating proteins by molecular weight can be one of the steps in two-dimensional gel electrophoresis. You can use the protein's molecular weight as a rough constraint to estimate whether it is reasonable to identify a particular protein in a certain fraction that was searched. |
| Pfam IDs | Displays the identification numbers of families of proteins. A special sequence comparison algorithm, called the Hidden Markov Model, groups proteins into families by comparing the sequences. Each family has its own ID number that starts with Pf ….<br>This column appears when the consensus workflow includes the Protein Annotation node. |
| Protein FDR Confidence | Displays the level of confidence of the identified protein groups as determined by the Protein FDR Validator node.<br>This column appears when the consensus workflow includes the Protein FDR Validator node. |
| Protein Group IDs | Displays the identification numbers of the referenced protein groups. |
| Score Sequest HT | Displays the protein score, which is the sum of the scores of the individual peptides.<br>This column appears when the consensus workflow includes the Protein Scorer node. |
| Sequence | Displays the sequence of amino acids that compose the peptide in the protein. |
| Sequence Coverage | Displays the parts of the protein sequence that were identified. The column ToolTip shows the number of amino acids composing the protein. You cannot export this column to a text file. |
| Sum PEP Score | Displays the scores that the Protein FDR Validator node calculates on the basis of the PEP values of the PSMs. The application uses these scores to rank the list of proteins. |
| Unique Sequence ID | Displays a unique identifier for the protein sequence. |
| WikiPathway Accessions | Displays the accessions from the Wiki Pathways database.<br>This column appears on the Proteins page when you include the Protein Annotation node in the consensus workflow. |
| WikiPathways | Displays the descriptions from the Wiki Pathways database. |

| | |
|---|---|
| | This column appears on the Proteins page when you include the Protein Annotation node in the consensus workflow. |
| Contaminant | Displays an X symbol next to the proteins marked as contaminants in the searched FASTA file or files.<br>This columns appears when the consensus workflow includes the Protein Marker node.. |
| Species Map | Extracts from the FASTA database the species names for proteins and displays and annotates them as colored entries in a distribution map.<br>This column appears only when you include the Protein Marker node in the consensus workflow and set its As Species Map parameter to True. |
| Species | Extracts from the FASTA database the species names for proteins and displays and annotates them as semicolon-separated text.<br>This column appears only when you include the Protein Marker node in the consensus workflow and set its As Species Names parameter to True. |

| Peptide Groups columns | |
|---|---|
| Parameters | Description |
| # Missed Cleavages | Displays the number of cleavage sites in a peptide sequence that a cleavage reagent (enzyme) did not cleave. This number excludes cases where an amino acid such as proline inhibits the cleaving enzyme. For example, if proline resides next to lysine or arginine, trypsin does not cleave the lysine or arginine. The application returns a 0 value when all peptides have been completely digested. |
| # Protein Groups | Displays the number of protein groups containing the peptide. |
| # Proteins | Displays the total number of identified proteins containing the peptide. |
| # PSMs | Displays the total number of identified PSMs in the peptide group, including those redundantly identified. |
| Abundance Ratios | Displays the abundance ratios as normal space values.<br>This column appears when there are sample ratios defined in the analysis setup. |
| Abundance Ratios (by Bio. Rep.) | Displays the abundance ratios of the biological replicates as normal space values. |
| Abundance Ratios (log2) | Displays the abundance ratios as log2 ratios. |
| Abundance Ratios (log2) (by Bio. Rep.) | Displays the abundance ratios of the biological replicates as log2 values. |
| Abundances | Displays the abundances of the samples before scaling and normalization. |

| | |
|---|---|
| Abundances (by Bio. Rep.) | Displays the abundance values of the biological replicates. |
| Abundances (by Bio. Rep.) Counts | Displays the number of abundance values used to calculate the abundances of the biological replicates. |
| Abundances (Grouped) | Displays the abundance values of the sample groups.<br>A grouped abundance value is calculated as the arithmetic mean of all the replicate abundance values within a sample group. You can specify the sample grouping on the Grouping and Quantification page when you set up an analysis.<br>This column appears when you group samples in the analysis setup and there is at least one sample group consisting of at least two samples. |
| Abundances (Grouped) Counts | Displays the number of samples with detected abundance values used to calculate the abundance of the whole sample group. |
| Abundances (Grouped) Standard Errors [%] | Displays the standard error of the abundance values of the samples in a sample group, normalized to the group's median abundance. |
| Abundances (Normalized) | Displays the normalized abundances values of the samples.<br>This column appears when you set the Normalization Mode parameter of the Precursor Ions Quantifier node or the Reporter Ions Quantifier node to Total Peptide Amount or Specific Protein Amount. |
| Abundances (Scaled) | Displays the normalized and scaled abundance values of the samples.<br>This column appears when you set the Scaling Mode parameter of the Precursor Ions Quantifier node or the Reporter Ions Quantifier node to On All Average or On Controls Average. |
| Abundances Counts | Displays the number of abundance values used to calculate the sample abundance. |
| Annotated Sequence | Displays the identified peptide sequence with the flanking residues from the master protein. |
| Charge (by Search Engine) | Displays the charge state of the peptide. |
| Checked | Indicates whether the item is selected. |
| Concatenated Rank (by Search Engine) | Displays the value for the "best" PSM found by the search engine. This value includes the rank of the decoy PSMs for the same spectra. |
| Confidence | Indicates a confidence level associated with the peptide group at the top level: green for high confidence, yellow for medium confidence, and red for low confidence. |

| | |
|---|---|
| Confidence (by Search Engine) | Indicates a confidence level associated with the peptide group at the top level: green for high confidence, yellow for medium confidence, and red for low confidence. |
| Found in Files | Displays the best confidence of the PSMs of the protein that the application identified in the files. The colors of the boxes represent the following:<br>•Green: high confidence<br>•Yellow: medium confidence<br>•Red: low confidence<br>Blue: found but unidentified PSM. Only the results from precursor<br>•ion quantification searches contain blue boxes.<br>This column appears only when you include the Data Distributions node in the consensus workflow and set its Show Found in Files parameter to True. |
| Found in Fractions | Displays the best confidence of the PSMs of the protein that the application identified in the fractions. The colors of the boxes represent the following:<br>•Green: high confidence<br>•Yellow: medium confidence<br>•Red: low confidence<br>Blue: found but unidentified PSM. Only the results from precursor<br>•ion quantification searches contain blue boxes.<br>This column appears only when you include the Data Distributions node in the consensus workflow and set its Show Found in Fractions parameter to True. |
| Found in Sample Groups | Displays the best confidence of the PSMs of the protein that the application identified in the sample groups. The colors of the boxes represent the following:<br>•Green: high confidence<br>•Yellow: medium confidence<br>•Red: low confidence<br>Blue: found but unidentified PSM. Only the results from precursor<br>•ion quantification searches contain blue boxes.<br>This column appears only when you include the Data Distributions node in the consensus workflow and set its Show Found in Sample Groups parameter to True. |
| Found in Samples | Displays the best confidence of the PSMs of the protein that the application identified in the samples. The colors of the boxes represent the following:<br>•Green: high confidence<br>•Yellow: medium confidence<br>•Red: low confidence<br>Blue: found but unidentified PSM. Only the results from precursor<br>•ion quantification searches contain blue boxes. |

| | |
|---|---|
| | This column appears only when you include the Data Distributions node in the consensus workflow and set its Show Found in Samples parameter to True. |
| m/z [Da] (by Search Engine) | Displays the mass-to-charge ratio of the precursor ion, in daltons. |
| Master Protein Accessions | Displays the unique identifiers (accessions) of all master proteins from all protein groups that include the given peptide sequence.<br>This column appears when you use the Protein Grouping node in the consensus workflow. |
| Master Protein Descriptions | Displays a description of the master proteins. |
| Modifications | Displays all the PTMs identified for the given peptide. The sites are displayed only for fixed modifications or for variable modifications when the confidence level is above the threshold set by the IMP-ptmRS node. They are indexed to the local sequence of the peptide, not to the master protein. |
| Modifications (All Possible Sites) | Displays all modifications found in the PSMs of a peptide group. They are indexed to the local sequence of the peptide, not to the master protein. |
| Modifications in Master Proteins | Displays the positions of the modifications in the master proteins only if the confidence level is above the threshold set by the IMP-ptmRS node. |
| Modifications in Master Proteins (All Sites) | Displays the positions of the modifications in the master proteins with confidence levels returned by the IMP-ptmRS node. |
| Percolator PEP (by Search Engine) | Displays the posterior error probability (PEP) of the best PSM of the peptide group that the search node identified.<br>This column appears when the consensus workflow includes the Percolator node. |
| Percolator q-Value (by Search Engine) | Displays the q-value of the best PSM of the peptide groups that the search node identified.<br>This column appears when the consensus workflow includes the Percolator node. |
| Percolator SVMScore (by Search Engine) | Displays the maximum Percolator score for the identified peptide group.<br>This column appears when the consensus workflow includes the Peptide Validator node. |
| Positions in Master Proteins | Displays the positions of the peptide in the master protein. |
| Protein Accessions | Displays the unique identifiers (accessions) of all proteins from all protein groups that include the given peptide sequence. |

| | |
|---|---|
| | The identifiers displayed in the Protein Group Accessions column are the same as those displayed in the Accession column on the Proteins and PSMs pages. |
| PSM Ambiguity | Displays the grouping status of the PSM in the peptide group. This column can display the following categories:<br>• Unambiguous: Indicates that this PSM is the only match that the application considered for this spectrum. There is no ambiguity that it needs to resolve.<br>• Selected: Indicates that the application selected this PSM from a set of two or more matches that it considered for the protein group inference process.<br>• Rejected: Indicates that the application rejected this PSM from a set of two or more matches that it considered for the protein group inference process.<br>• Ambiguous: Indicates that the application considered two or more peptide matches to the same spectrum for the protein group inference process. If you set the Apply Strict Maximum Parsimony Principle parameter of the Protein Grouping node to True in the consensus workflow, the application resolves this ambiguity to Selected or Rejected.<br>• Unconsidered: Indicates that the application did not consider this PSM for the protein group inference process because it did not meet the requirements specified in the Protein Grouping (Enabled) settings in the Result Filters page. |
| Quan Info | Indicates why the application used or did not use a peptide in quantification:<br>• No Quan Values: Indicates that there are no usable quantification results.<br>(For reporter ion quantification)<br>• No Quan Labels: A label did not modify the peptide.<br>(For precursor ion quantification)<br>• Inconsistently Labeled: Peptides do not always display the expected amount of isotopic labeling.<br>• Indistinguishable Channels: The peptide does not have amino acids that can have the specified labels, or it has the specified labels but has defined channels with the same delta masses, rendering the channels indistinguishable.<br>• Incompatible Labels: Indicates that the composition of a modification is not known.<br>• Redundant: Indicates that the quantification results of this peptide group are redundantly used.<br>• Is Not Reliable: Indicates that all PSMs classified as A1 triggers are excluded from the quantification. |

| | |
|---|---|
| | • Excluded Modification: Indicates that a peptide group has specified modifications, which are defined by several parameters of the Precursor Ions Quantification node.<br>• Shared: Indicates that the peptide group is usable for quantification but is shared among different protein groups or proteins, depending on the setting of the Consider Protein Groups for Peptide Uniqueness parameter in the Reporter Ions Quantifier node. |
| Quality PEP | Displays the posterior error probability (PEP) for the identified peptide group.<br>This column appears when the consensus workflow includes the Peptide Validator node. |
| Quality q-value | Displays the q-value for the identified peptide group.<br>This column appears when the consensus workflow includes the Peptide Validator node. |
| Rank (by Search Engine) | Displays the ordering of peptides by rank. |
| RT [min] (by Search Engine) | Displays the retention time when the peptide was observed, in minutes. |
| Search Engine Rank (by Search Engine) | Displays the original rank assigned by the search engine for all PSMs and peptide groups. |
| Sequence | Displays the sequence of amino acids that compose the peptide in the peptide group. |
| Sequence Length | Displays the number of amino acid in a sequence. |
| SVM_Score | Displays the maximum Percolator score for the identified peptide group. |
| Theo. MH+ [Da] | Displays the mass calculated from the sequence, the amino acid masses, and the defined modification masses, as opposed to the measured mass. |
| ΔCn (by Search Engine) | Displays the normalized score difference between the currently selected PSM and the highest-scoring PSM for that spectrum. |
| ΔM [ppm] (by Search Engine) | Displays the difference between the theoretical mass of the peptide and the experimental mass of the precursor ion, in parts per million. |
| Δm/z [Da] (by Search Engine) | Displays the difference between the measured charged mass (m/z [Da]) and the theoretical mass of the same charge (z). |
| ΔScore (by Search Engine) | Displays a measure of the difference between the top two scores for the peptides identified by that spectrum. The application calculates this score as ΔScore = Score(Rank N Peptide) – Score(Rank 1 Peptide)/Score(Rank 1 Peptide). |
| Contaminant | Displays an X symbol next to the proteins marked as contaminants in the searched FASTA file or files. |

| | |
|---|---|
| | This column appears when the consensus workflow includes the Protein Marker node. |
| Ratio columns | Display the corrected ratio of the intensity of the reporter ion in a sample to the intensity of the reporter ion in the control sample. |
| Sequences in Proteins | Displays the flanking residues as a consensus sequence in square brackets; if multiple proteins share the peptide, this column lists all the different flanking residues.<br>This column appears when you use the Peptide in Protein Annotation node in the consensus workflow. |
| XCorr search_engine_number node_number | Scores the number of fragment ions that are common to two different peptides with the same precursor mass and calculates the cross-correlation score for all candidate peptides queried from the database (Sequest HT searches only). |
| Protein Quan Usage | Shows which peptide abundances are used for protein quantification.<br>•Used: Indicates that the node used quantification abundances.<br>Not Used: Indicates that the node did not use quantification<br>•abundances. |
| Species Map | Extracts from the FASTA database the species names for proteins and displays and annotates them as colored entries in a distribution map.<br>This column appears only when you include the Protein Marker node in the consensus workflow and set its As Species Map parameter to True. |
| Species | Extracts from the FASTA database the species names for proteins and displays and annotates them as semicolon-separated text.<br>This column appears only when you include the Protein Marker node in the consensus workflow and set its As Species Names parameter to True. |
| Areas | Displays a graphical distribution map showing the distribution of areas over the specified sample groups specified on the Grouping & Quantification page of the Study Definition page<br>This column appears only when you include the Precursor Ions Quantifier or the Reporter Ions Quantifier node in the processing workflow and also the Data Distributions node in the consensus workflow. |
| P-Value | Displays the p value for the sample group calculated by running the Tukey HSD test (post hoc) after an analysis of variance (ANOVA) test. The p value is a number between 0 and 1. If the null hypothesis states that there is no difference between the sample groups for the variance tests: |

| | |
|---|---|
| | • A low P value means there is a low probability that the null hypothesis is false; that is, the sample groups show a significant difference.<br>• A high P value means there is a high probability that the null hypothesis is true; that is, the sample groups do not show a significant difference. |
| Adj. P-Values | Displays the p-values adjusted by using the Benjamini-Hochberg correction for the false discovery rate. |
